# Just Add Structure: Protein Language Models Combined with Structural Equivariance Excel at Protein Tasks

**DOI:** 10.64898/2026.05.28.728196

**Authors:** Qurat-ul-ain, Carlos Outeiral, Matteo Cagiada, Yee Whye Teh, Charlotte M. Deane

## Abstract

Accurate *in silico* prediction of protein properties, functional fitness, and mutational effects remains a central challenge in protein engineering and therapeutic design. While Protein Language Models (PLMs) successfully capture rich evolutionary and functional constraints from sequence data, they only indirectly encode the spatial and geometric information that fundamentally governs protein function. Consequently, state-of-the-art approaches typically rely on extensive fine-tuning, ensembling, or the incorporation of handcrafted structural features to achieve competitive accuracy, making them computationally expensive and difficult to scale. In this work, we demonstrate that explicit geometric modeling can substitute for, and in most cases outperform, large-scale PLM fine-tuning, with much higher parameter efficiency. Our approach, ProtEGNN, pairs PLM residue representations with a lightweight ***E*(3)**-Equivariant Graph Neural Network, competing with or achieving state-of-the-art performance across eight different benchmarks in protein property, mutational effect and function prediction, while needing 100–1000***×*** fewer parameters than competing methods. Even when protein structure is combined with representations from ESM2-T6, a small 8M-parameter PLM, ProtEGNN matches fine-tuned sequence-only approaches based on substantially larger PLM backbones, while training orders of magnitude fewer parameters. Together, these results highlight geometric inductive bias as a powerful and scalable alternative to task-specific fine-tuning of large PLMs for protein modeling.

## 1 Main

Protein engineering is a cornerstone of modern biotechnology, enabling the design and optimisation of proteins with desirable functions and properties for applications in medicine, industry, and synthetic biology. Central to this effort is the challenge of understanding how amino acid sequence gives rise to protein function and biophysical behaviour, with the goal of tailoring proteins for improved stability [1], solubility [2], catalytic performance [3], binding, and other therapeutically relevant traits. Traditional approaches, including rational design and directed evolution, have delivered major advances [4], but remain fundamentally constrained by the vastness of protein sequence space and the labour-intensive cost of experimental validation [5]. These limitations have become even more apparent as recent advances in protein design [6–8] have expanded our ability to generate novel protein backbones and binding scaffolds, shifting the field’s central challenge from design itself towards the optimisation of early candidates into viable therapeutic and industrial products.

A major bottleneck in this optimisation process is the experimental determination of protein properties, which remains slow, expensive, and labour-intensive. Measurements typically require recombinant protein expression and purification, often taking several weeks and costing thousands of dollars, followed by careful sample preparation, data collection, and interpretation. Computational approaches for analysing protein properties *in silico* are therefore essential for accelerating protein engineering, as they reduce dependence on low-throughput experimental screening and help alleviate key bottlenecks in drug discovery and synthetic biology.

The most common tools used for protein analysis are Protein Language Models (PLMs) trained on large-scale sequence databases, which achieve strong performance in tasks such as property prediction, mutational-effect estimation and protein engineering [9–12]. Models such as the ESM suite (ESM2, ESM3, ESMc) [13, 14], ProtTrans [15], Ankh3 [16] and others [17, 18] capture evolutionary and functional constraints from protein sequences and have even been shown to implicitly reflect aspects of the tertiary structure, including fold classes, residue–residue contact patterns and molecular motions [19–21]. However, since PLMs are trained only on amino acid sequences, this structural knowledge is indirect: it emerges from statistical regularities in sequence co-variation rather than symmetry-aware geometric reasoning. This distinction is important because many protein properties are fundamentally governed by spatial interactions [22]. For example, solubility depends on surface exposure and charge distribution, and thermodynamic stability depends on core packing and hydrogen-bond networks. Numerous studies suggest that structural data [12, 23] can inform protein properties in ways that sequences alone cannot, as the three-dimensional configuration of a protein is a major determinant of its biophysical and functional characteristics [24, 25]. Improving how deep learning models incorporate structural information therefore represents a clear route to more accurate prediction of protein properties and function.

However, rather than focusing on principled integration of protein geometry, the predominant strategy used by state-of-the-art models for protein tasks has been to apply increasingly sophisticated fine-tuning pipelines [26] and parameter-efficient adaptation methods on ever-larger PLMs [27]. Many of these approaches further rely on ensembling [2] or mix-and-match frameworks that combine embeddings from multiple PLMs with diverse downstream architectures [28, 29], often requiring thousands of experimental configurations to achieve competitive performance [30].

Although these strategies can be effective, they also introduce substantial practical and conceptual limitations. Fine-tuning can lead to catastrophic forgetting [31], whereby specialization to a particular downstream task degrades the broader representations acquired during pretraining and can result in worse performance than the original frozen model on other tasks [32]. At the same time, the rapid growth of PLMs, from roughly 10^6^ to more than 10^9^ parameters, has made task-specific adaptation of these models increasingly difficult and computationally expensive. As a result, current approaches often come with growing engineering complexity, higher training costs, and greater challenges for reproducibility.

Moreover, the emphasis on elaborate adaptation of PLMs treats sequence representations as the dominant, and often sufficient, lever for improvement, while principled geometric modeling remains underutilized. Even when structure is incorporated into PLMs, it is often done indirectly through handcrafted features, coarse distance bins, or quantazied 3D token representations that are appended to sequence embeddings rather than modeled as continuous 3D geometry [33, 34]. As a result, many methods are costly to train and adapt yet still lack explicit structural inductive biases. We challenge this fine-tune-first approach and contend that joint modeling of static PLMs representations with explicit protein structure using Equivariant Graph Neural Networks (EGNNs) [35] is a direct and efficient path to improved performance in protein tasks.

In this work, we demonstrate that structural geometry, when modeled explicitly, can rival the predictive power of massive PLMs while using 100–1000× fewer trainable parameters, rendering large-scale fine-tuning unnecessary. We introduce ProtEGNN, a simple framework for integrating explicit structural information with PLMs, and show that it is competitive with leading approaches across a diverse set of protein prediction tasks despite being substantially simpler and far smaller in scale. ProtEGNN (Fig 1) represents each protein as a graph, where nodes correspond to the 3D coordinates of backbone C_*α*_ atoms in a predicted structure. Each node is initialized with a feature vector given by embeddings from the final layer of a pretrained PLM; these embeddings are extracted once and treated as fixed inputs during training. Edges connect spatially proximal residues, and an EGNN updates the node features in an equivariant manner, enabling the model to combine sequence-derived information with explicit structural geometry.

**Fig. 1.**
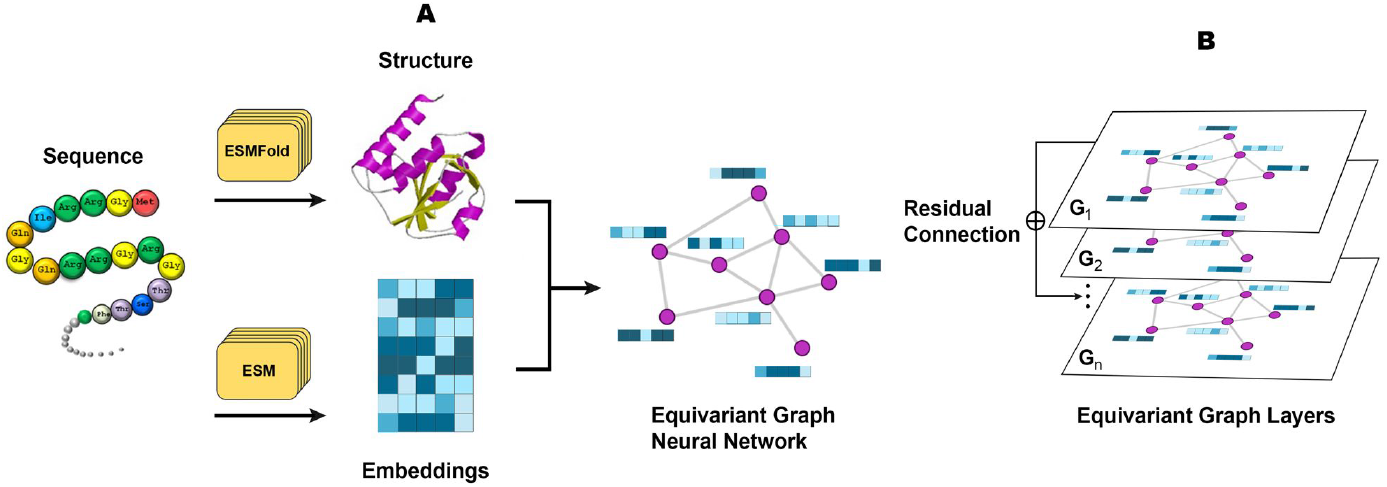
Integration of protein sequence and structure data using a EGNN architecture in ProtEGNN. **Panel A**: a protein sequence is processed through two distinct pathways: ESMFold predicts the 3D structure of the protein, while ESM (ESMc or ESM2-T6) generates sequence embeddings. The predicted structure and sequence embeddings are then combined to construct a graph representation, where nodes correspond to amino acids and edges represent spatial relationships. **Panel B**: the graph is passed through multiple EGNN layers. Residual connections are used between layers to maintain information flow and improve learning stability.

We evaluated whether incorporating explicit protein structure can compensate for, or reduce the need for, adaptation of PLMs and established a baseline by comparing ProtEGNN to the pretrained and fine-tuned version of the same PLM, isolating the effects of model scale, fine-tuning, and explicit geometry. Moreover, across eight datasets spanning protein property prediction, mutational fitness prediction, and protein function annotation, ProtEGNN consistently matches or outperforms the leading methods in protein tasks, including models with far greater architectural complexity and 100–1000× more parameters. In particular, ProtEGNN achieves substantial improvements on solubility and thermostability prediction, establishing new state-of-the-art performance on both tasks by substantial margins. Through experiments in mutational effect prediction (GB1, GFP, SBI), we show that modeling protein structure with an *E*(3)-equivariant graph, even when paired with a small PLM (ESM2-T6; 8M parameters) backbone and a *single wild-type structure*, delivers gains that rival state-of-the-art methods relying on large-scale PLMs and task-specific fine-tuning. These results demonstrate that principled geometric inductive biases can be a more effective lever for performance than additional model scale or fine-tuning.

## 2 Results

We assessed our method, ProtEGNN (Figure 1), on eight datasets spanning three protein modelling regimes: protein property prediction, mutational-effect prediction, and protein function annotation. These tasks differ both in the biological questions they address and in the level at which predictions are made. Protein property prediction and function annotation are formulated as graph-level tasks, in which the goal is to infer a single label or score for an entire protein from its sequence and structure, whereas mutational-effect prediction is treated as a node-level setting, where predictions depend on the local consequences of residue substitutions within a fixed structural context. Across these benchmarks, the outputs include regression, binary classification, and multiclass classification tasks, which we evaluate using Spearman correlation (SPR), accuracy (ACC), and Receiver Operating Characteristic Area Under the Curve (ROC AUC), respectively. Together, these datasets capture a broad range of biological settings, from sequence-diverse proteins requiring global generalization to mutation-centric landscapes requiring sensitivity to local structural constraints. This diversity provides a stringent test of whether structure-aware contextualization of sequence information through using equivariant graphs offers consistent value across distinct classes of protein prediction problems.

### 2.1 Baseline Experiments

Our goal with ProtEGNN is to show that joint modeling of explicit 3D protein structure with PLM representations is an effective and efficient strategy for protein tasks, and should be prioritized before resorting to costly fine-tuning of large PLMs. We instantiated ProtEGNN with representations from three pretrained PLMs spanning a wide range of scales: the small ESM2-T6 model with 8M parameters [13], the medium-scale ESMc-M model with 600M parameters [14], and the larger Ankh-large model with 1.2B parameters [36]. To contextualize the performance of ProtEGNN and isolate the contribution of explicit geometry, we evaluate three settings for each PLM backbone: a pretrained PLM used as a static feature extractor, a task-adapted PLM, and ProtEGNN, which augments fixed PLM residue embeddings with protein structure.

We summarize the effect of explicit geometric modeling across tasks and PLM scales (Table 1). See the Methods section for a full description of all datasets and tasks. For the small ESM2-T6 backbone, ProtEGNN yields consistent gains over both pretrained and fine-tuned baselines across every benchmark, demonstrating that explicit geometry can substantially enhance the predictive power of even compact sequence representations. For the ESMc-M backbone, ProtEGNN achieves the best performance on five out of six tasks, with the only exception being Stability, where full fine-tuning performs better. Ankh-large provides an additional test of whether the benefits of explicit geometric modeling persist when the underlying sequence model is already very large. ProtEGNN improves over the pretrained baseline across all tasks and outperforms the fine-tuned model on Meltome, Stability, Solubility, and subcellular localization, while matching the performance on GFP and remaining below fine-tuning on GB1. These results suggest that incorporating structure through an equivariant graph head improves or matches performance over pretrained sequence-only baselines across most tasks regardless of PLM backbone size. The following sections compare ProtEGNN against leading methods on each task, and contextualize these gains by comparing the number of trainable parameters required to achieve them.

**Table 1.**
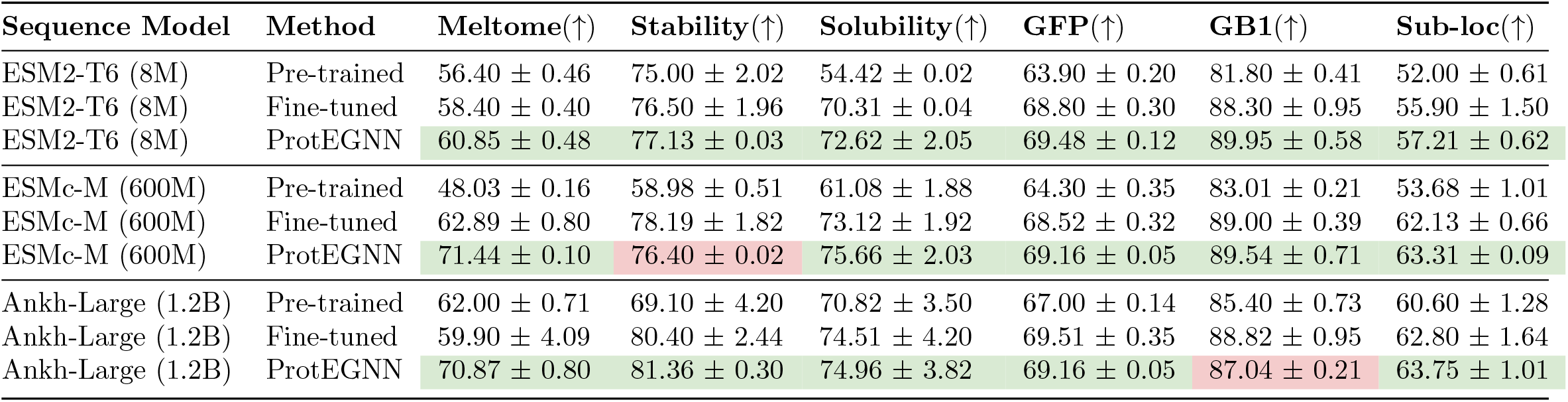
Effect of PLM scale and explicit geometric modeling across tasks. For ESM2-T6 and Ankh-Large, pre-trained and fine-tuned results are as reported in [27], except for solubility, which we evaluate in this work using similar protocol. Green cells highlight ProtEGNN results that match or exceed the fine-tuned model; red cells highlight ProtEGNN results that fall below that fine-tuned counterpart.

### 2.2 Protein Property Prediction

#### 2.2.1 Solubility

We use the PSI Biology solubility dataset [37], curated and cleaned by [2], containing 11,226 *E. coli* –expressed proteins labeled as soluble or insoluble. We compare our method, ProtEGNN, against Prot-T5 [15], a large 1.2B parameter protein language model, ESM-MSA [38], which leverages Multiple Sequence Alignments and NetSolP [2], a previous state-of-the-art solubility predictor specifically trained and tuned on the PSI-Biology dataset (Table 2). NetSolP relies on both fine-tuning and ensembling of multiple ESM models [19]. PSI-Biology results for Prot-T5, ESM-MSA, and NetSolP are as reported in [2]. Despite being orders of magnitude smaller, both ProtEGNN variants perform competitively; with ProtEGNN(ESMc-L) setting the new state-of-the-art while using 100x less parameters.

**Table 2.** Solubility prediction performance and model size comparison. Results are shown in ascending order of performance (ROC-AUC). Higher is better. Bold values indicate our method.

| PSI-Biology (5-fold CV) |  |  | Price (zero-shot) |  |  |
| --- | --- | --- | --- | --- | --- |
| Method | Params | ROC-AUC(↑) | Method | Params | ROC-AUC(↑) |
| Prot-T5 | 1.2B | 0.73 | ProtSolM | 3.2M | 0.55 |
| NetSolP | 650M | 0.73 | PLMSoL | 7.3M | 0.60 |
| **ProtEGNN (t6)** | **500K** | **0.73** | Prot-T5 | 1.2B | 0.73 |
| ESM-MSA | 100M | 0.75 | **ProtEGNN (t6)** | **500K** | **0.74** |
| **ProtEGNN (ESMc-L)** | **1.2M** | **0.78** | ESM-MSA | 100M | 0.75 |
|  |  |  | NetSolP | 650M | 0.76 |
|  |  |  | **ProtEGNN (ESMc-L)** | **1.2M** | **0.77** |

To assess out-of-distribution performance, we tested leading solubility predictors on the Price solubility dataset [39], which was not used in training ProtEGNN (Table 2). ProtSolM [40] integrates an EGNN directly with a PLM, together with handcrafted physicochemical structural descriptors, and performs end-to-end training with gradients propagated through the entire combined architecture. Similarly, PLMSol [28] aggregates embeddings from multiple large PLMs and couples them with an ensemble of classifiers (e.g. MLPs, Light Attention, CNN–BiLSTM), resulting in a resource-intensive approach. ProtEGNN(ESMc-L) achieved the best zero-shot performance with ROC AUC of 0.77 using only 1.2M parameters, outperforming substantially larger models. Even our smaller variant, ProtEGNN(t6), remained competitive (0.74, 500K), surpassing ProtSolM (0.55, 3.2M) and PLMSoL (0.60, 7.3M).

#### 2.2.2 Stability

On the first thermostability benchmark, Meltome, [41] which contains ~ 23,300 protein sequences with melting temperatures (*T*_*m*_), we observe that PLM capacity plays a larger role than in other tasks. ProtEGNN(t6) variant achieves only 0.61 SPR, trailing substantially behind approaches built on large pretrained backbones (Table 3). Scaling the PLM size within our framework yields a large jump: ProtEGNN(ESMc-L) reaches 0.79 SPR, outperforming all compared methods by a significant margin. Prot-T5-FT [27] attains strong performance after adapting only 3.5M out of the 1.2B parameters of Prot-T5 [15], yet lacks structural bias. Other baselines rely on resource-intensive pretraining and tuning pipelines: PLM-Fit [30] benchmarks combinations of multiple PLMs with methods such as feature extraction and bottleneck adapters across five protein engineering datasets, which required 3,000+ experiments and PRIME [26] is pre-trained on 96M bacterial sequences annotated with optimal growth temperatures and then fine-tuned on downstream tasks.

**Table 3.**
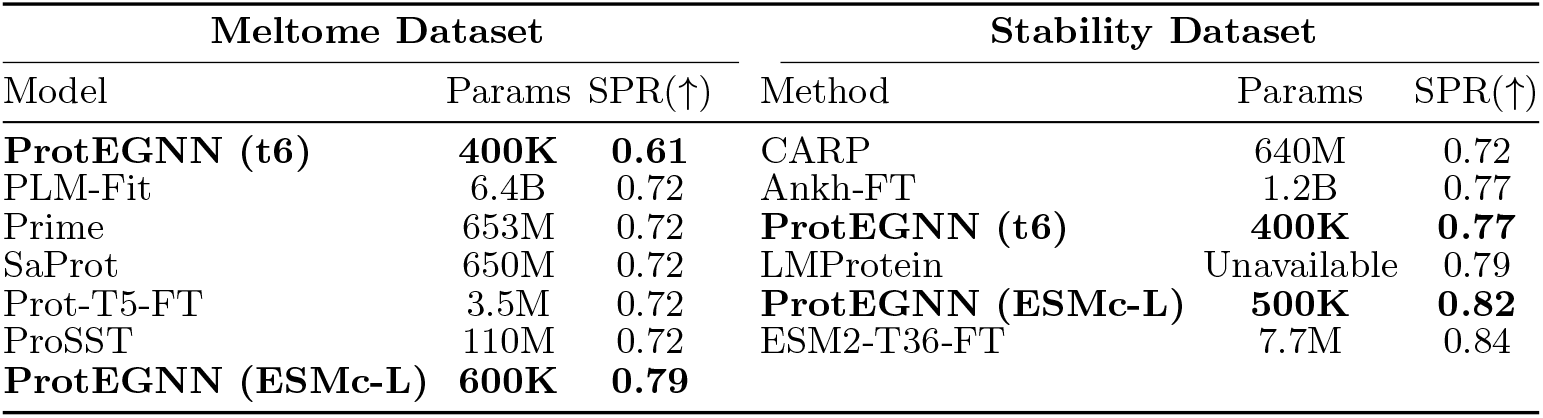
Performance and model size comparison on Meltome and Stability datasets. Results are shown in ascending order of performance (SPR). Higher is better. Bold values indicate our method.

| Meltome Dataset |  |  | Stability Dataset |  |  |
| --- | --- | --- | --- | --- | --- |
| Model | Params | SPR(↑) | Method | Params | SPR(↑) |
| **ProtEGNN (t6)** | **400K** | **0.61** | CARP | 640M | 0.72 |
| PLM-Fit | 6.4B | 0.72 | Ankh-FT | 1.2B | 0.77 |
| Prime | 653M | 0.72 | **ProtEGNN (t6)** | **400K** | **0.77** |
| SaProt | 650M | 0.72 | LMProtein | Unavailable | 0.79 |
| Prot-T5-FT | 3.5M | 0.72 | **ProtEGNN (ESMc-L)** | **500K** | **0.82** |
| ProSST | 110M | 0.72 | ESM2-T36-FT | 7.7M | 0.84 |
| **ProtEGNN (ESMc-L)** | **600K** | **0.79** |  |  |  |

Several methods additionally incorporate structural signals: ProSST [34] learnt a structure-quantized representation with disentangled sequence–structure attention, and SaProt [33] augmented PLMs with quantized 3D structural tokens to inject geometry directly into the sequence model. In comparison, our approach keeps the PLM frozen and uses its embeddings as node representations for an EGNN, achieving state-of-the-art SPR with orders-of-magnitude fewer parameters than millioins-to-billions parameter PLM based pipelines. Overall, the relatively weak performance of ProtEGNN(t6), paired with the clear gains from ProtEGNN(ESMc-L), suggests that thermostability prediction depends heavily on rich pretrained representations and that the way structure is integrated into PLMs is consequential, motivating a shift from scale-first to geometry-aware design.

On the second stability benchmark [42], which measures protease susceptibility for de novo–designed mini-proteins, a different pattern emerges (Table 3). In contrast to Meltome, the small-backbone variant ProtEGNN(t6) achieves strong performance (SPR = 0.77), matching Ankh-FT and outperforming sequence-only baselines such as CARP [43]. This suggests that for this dataset—where generalization is evaluated locally around optimized designs—explicit geometric modeling can compensate for limited PLM capacity. Scaling to ProtEGNN(ESMc-L) further improves performance to 0.82 SPR, surpassing CARP (0.72) [43] which combined PLMs with Convolutional architecture, and LMProtein (0.79) [29], a hybrid architecture that stacks CNNs, LSTMs, and MLPs on top of ESM-2 embeddings. The LoRA finetuned ESM2-T36-FT [27] achieves the highest score (0.84), albeit using orders-of-magnitude more parameters.

Together with the Meltome results, this comparison indicates that thermostability prediction benefits from both rich pretrained representations and explicit structure, but that the relative importance of PLM scale versus geometry depends on the nature of the stability task; global across diverse proteins versus local around optimized designs.

### 2.3 Mutational Effect Prediction

For each dataset, we use a *single wild-type structure* to represent all variants, fixing the underlying geometry across the mutational landscape. Even without modeling the explicit mutation-induced structural rearrangements, our equivariant geometric model captures local physical constraints that remain informative under sequence perturbations, enabling state-of-the-art mutational effect prediction.

#### 2.3.1 GFP

We report results on the GFP [44] dataset, a regression task where ~54,000 sequences are mapped to their log-fluorescence intensity. ProtEGNN(T6) achieves state-of-the-art performance (SPR = 0.70), equal to Ankh-FT model [27], while training only 500K task-specific parameters on top of a small 8M-parameter PLM backbone (Table 4). Notably, Ankh-FT applied LoRA on top of the 1.9B parameter Ankh backbone, resulting in millions of trainable parameters. ProtEGNN achieves the same performance using fewer parameters (4.9M vs 500K) and much smaller PLM backbone (1.9B vs 8M).

**Table 4.**
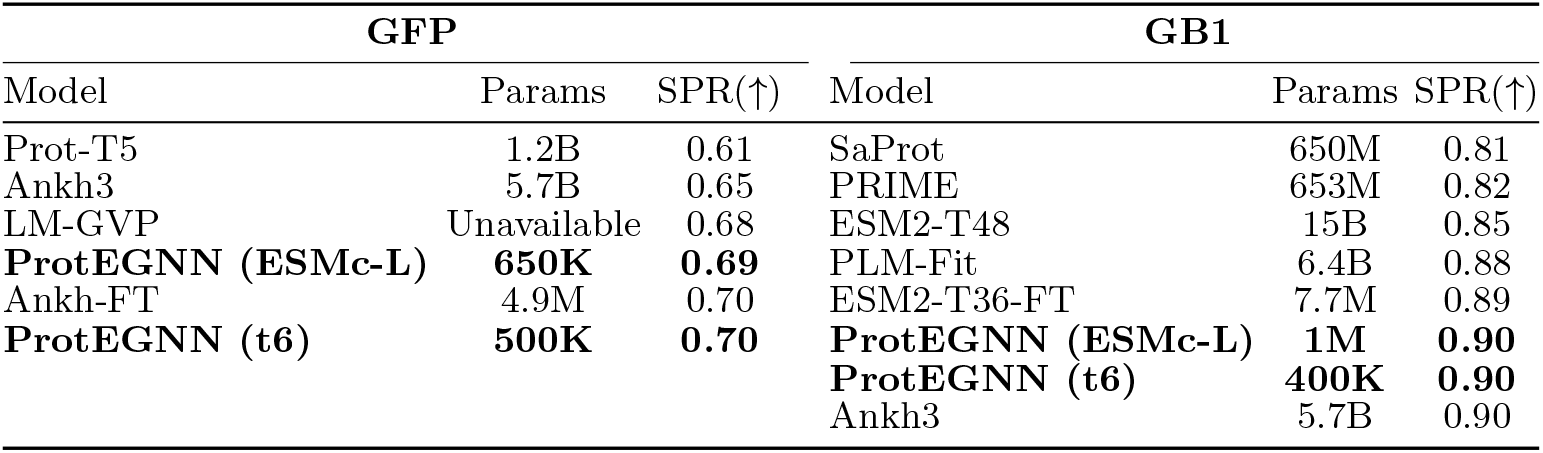
GFP and GB1 mutational effect prediction performance and model size comparison. Results are shown in ascending order of performance (SPR). Higher is better. Bold values indicate our method.

| GFP |  |  | GB1 |  |  |
| --- | --- | --- | --- | --- | --- |
| Model | Params | SPR(↑) | Model | Params | SPR(↑) |
| Prot-T5 | 1.2B | 0.61 | SaProt | 650M | 0.81 |
| Ankh3 | 5.7B | 0.65 | PRIME | 653M | 0.82 |
| LM-GVP | Unavailable | 0.68 | ESM2-T48 | 15B | 0.85 |
| **ProtEGNN (ESMc-L)** | **650K** | **0.69** | PLM-Fit | 6.4B | 0.88 |
| Ankh-FT | 4.9M | 0.70 | ESM2-T36-FT | 7.7M | 0.89 |
| **ProtEGNN (t6)** | **500K** | **0.70** | **ProtEGNN (ESMc-L)** | **1M** | **0.90** |
|  |  |  | **ProtEGNN (t6)** | **400K** | **0.90** |
|  |  |  | Ankh3 | 5.7B | 0.90 |

Both ProtEGNN variants outperform substantially larger models such as Prot-T5 (1.2B parameters) [15] and Ankh3 (5.7B parameters) [16]. ProtEGNN also performs better than LM-GVP [45] (0.68), which stacked an EGNN in front of a PLM [15] and jointly trained the models, modifying and fine-tuning the PLM embeddings.

#### 2.3.2 GB1

We observe a similar pattern in the GB1 [41] binding landscape (Table 4), which measures binding affinity of ~8,700 variants of the immunoglobulin-binding domain of Protein G. Both ProtEGNN(t6) and ProtEGNN(ESMc-L) reach an SPR of 0.90, matching the best-performing 5.7B parameter Ankh3 model [16]. This parity is achieved with as few as 400K trainable parameters in ProtEGNN(t6). ESM2-T36-FT, as reported in [27] applies LoRA to a 3B parameter PLM backbone, reducing the number of trainable parameters to only 7.7M; however, despite this adaptation, it still underperforms ProtEGNN, suggesting the limitations of parameter-efficient fine-tuning when structural inductive biases are absent. Crucially, our results show that adding a single wildtype structure to even the smallest 8M parameter backbone delivers predictive power equivalent to multi-billion parameter sequence models without the need for extensive fine-tuning or structure predictions for every variant.

#### 2.3.3 Stable-But-Inactive Variant Prediction

Mutations that impair protein function can do so through distinct mechanisms. Some act indirectly by destabilizing the native fold, whereas others, including stable-but-inactive (SBI) mutations, preserve overall stability but directly disrupt catalytic residues, molecular interactions, or allosteric regulation. Most mutation-effect predictors, however, are designed only to identify deleterious variants and do not distinguish between the molecular mechanisms underlying functional loss. We therefore use SBI prediction as a test of whether explicit structural modelling enables a model to move beyond generic deleteriousness and separate direct functional perturbation from structural destabilization.

Fig 2 shows that ProtEGNN based on ESMc-S, 300M PLM backbone, accurately predicts SBI variants across the eight test systems, demonstrating that structural information is necessary to separate unstable mutations from variants that are inactive despite preserving fold stability. This is a strict test of mechanistic understanding: rather than simply detecting deleteriousness, the model must distinguish direct disruption of function from indirect loss of function through destabilization. Across both ROC AUC and AUPRC, ProtEGNN outperforms competing baselines, including other protein language models (ESM-1b) [13], structure-based predictors (ESM-IF) [46], and dedicated SBI detection methods such as FuncESM [47].

**Fig. 2.**
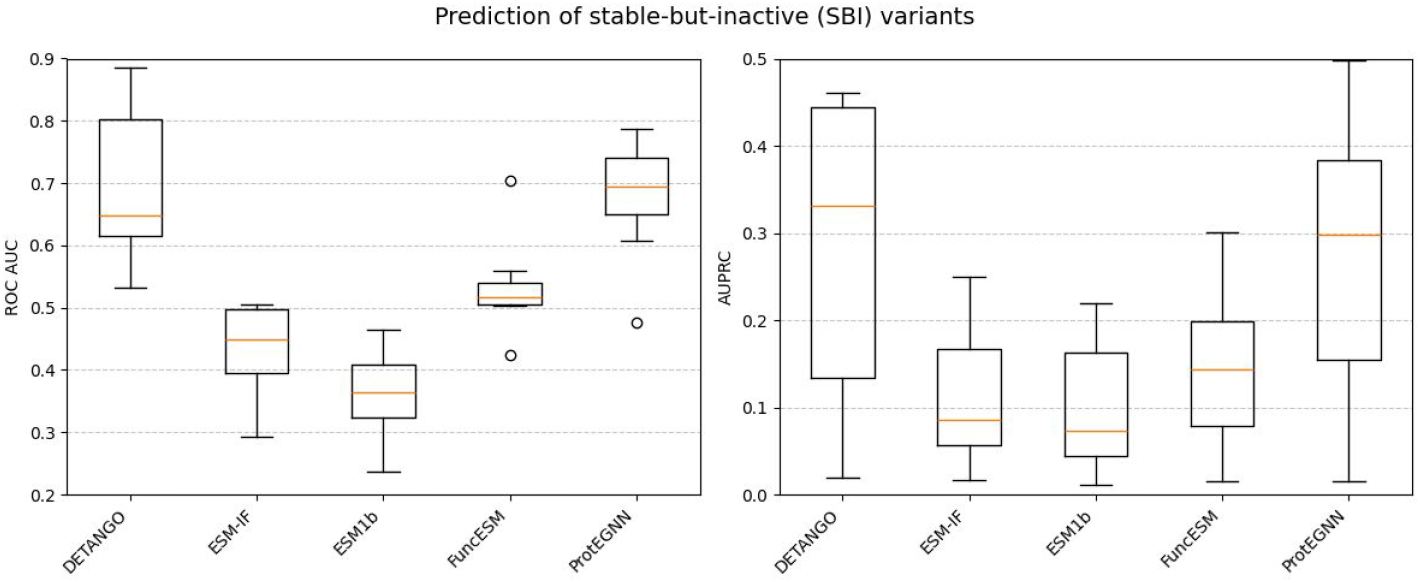
Performance comparison across five methods on SBI classification, evaluated using ROC AUC (left) and AUPRC (right). Boxplots summarize performance across 8 test proteins. ProtEGNN (ProteoGNN) consistently outperforms sequence-only baselines and remains competitive with DETANGO, a specialised method for SBI prediction.

ProtEGNN achieves performance comparable to DETANGO [48], the current state-of-the-art method for SBI detection, despite using far less supervision and structural input. DETANGO is pretrained on millions of protein sequences, whereas ProtEGNN is trained using DMS assays from only three proteins and relies on a single wild-type structure per protein. These findings show that explicit modelling of the wild-type fold allows the model to reason about mutational consequences beyond generic pathogenicity, enabling discrimination between stable but functionally impaired variants and those that act through destabilization.

### 2.4 Protein Function Annotation

#### 2.4.1 Subcellular Localization

On the subcellular localization benchmark [49], a 10-class per-protein classification task, ProtEGNN achieves competitive accuracy with orders-of-magnitude fewer parameters than large PLM baselines (Table 5). In particular, ProtEGNN matches the performance of Prosst-T5 [32] (ACC = 0.57) while using only 400K parameters. Even though Prosst, a fine-tuned derivative of ProtT5 [15], incorporates structural information, its reduced performance relative to ProtT5 and LA-Prot-T5 [10] (0.57 vs. 0.65) highlights the effect of task specialization through fine-tuning, where adapting a PLM for one objective can degrade its performance on others.

**Table 5.**
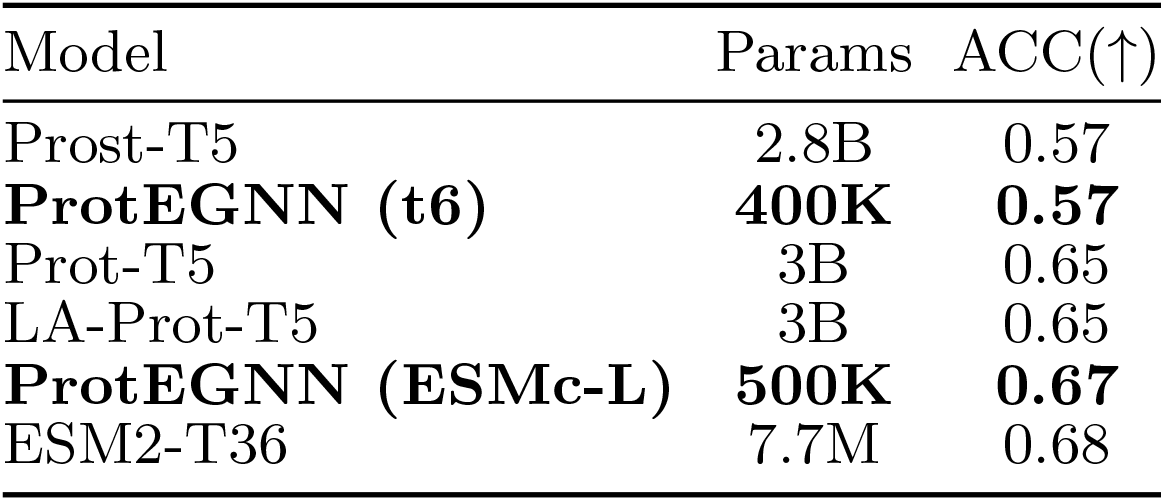
Subcellular Localization performance and model size comparison. Results are shown in ascending order of performance (ACC).

In contrast, ProtEGNN(ESMc-L) attains an accuracy of 0.67 with just 500K parameters, closely approaching the performance of the fine-tuned ESM2-T36 (0.68) reported in [27]. It is important to note that Schmirler et al. [27] employed parameter efficient fine tuning techniques to reduce the trainable parameters of ESM2-T36 from 3B to 7.7M. However, even parameter-efficient fine-tuning techniques approaches incur significantly higher parameters compared to our method, which simply adds a lightweight geometric head to frozen PLM representations. These results reinforce the idea that adding explicit structural inductive biases is a more robust and efficient alternative to fine-tuning large PLMs, even when parameter-efficient techniques are used.

### 2.5 Discussion

Our experiments convey a clear and actionable message: explicit, equivariant geometric modeling can substitute for, and in most cases outperform, large-scale PLM fine-tuning, while being 100-1000× more parameter-efficient. By pairing frozen PLM residue embeddings with a lightweight *E*(3)-equivariant GNN, ProtEGNN achieves strong performance across solubility, thermostability, and mutational-effect prediction while training fewer than 1M task-specific parameters. Our findings expose a broader limitation of current practice: fine-tuning large PLMs for each downstream task is not a viable long-term strategy. Beyond the computational burden, extensive fine-tuning sometimes leads to task specialization at the cost of generality, as seen in the case of ProSST, where subcellular localization performance degrades after fine-tuning. Our results on mutational landscapes demonstrate that geometric deep learning offers distinct advantages for protein modelling, enabling lightweight models to capture local physical constraints and recover much of the predictive benefit often attributed to massive, fine-tuned PLMs. This trend extends even to the more mechanistically demanding SBI prediction setting, where ProtEGNN achieves performance comparable to DETANGO despite relying only on a single wild-type structure per protein and far less large-scale pretraining, highlighting that explicit structural context can provide information that scale alone does not.

The design of ProtEGNN offers several practical advantages. First, it is highly parameter-efficient, delivering competitive performance while using around 100× fewer trainable parameters than approaches based on adapting billion-parameter PLMs or assembling large ensemble pipelines, thereby substantially reducing training, tuning, replication costs, and memory requirements. Second, it is also data-efficient: in the SBI prediction setting, ProtEGNN achieves performance comparable to specialised large-scale methods despite being trained on only a small number of proteins and using just a single wild-type structure per protein. Third, it provides a principled mechanism for injecting structural inductive biases, namely equivariance and continuous three-dimensional geometry, that are difficult to recover through fine-tuning alone and appear particularly valuable on benchmarks with genuine structural diversity, such as solubility and thermostability. More broadly, our findings underscore the importance of appropriate inductive biases in model design, and suggest that higher-fidelity representations and biology-aware architectures may yield greater gains in protein modelling than parameter scaling or increasingly elaborate adaptation of protein language models.

At the same time, our results highlight important task-dependent limits. The performance gap between ProtEGNN(T6) and ProtEGNN(ESMc-L) on thermostability and subcellular localization, compared to ProtEGNN(T6) matching state-of-the-art accuracy on mutational fitness tasks (GFP, GB1), indicates that PLM capacity matters more for some tasks than others. In particular, small frozen embeddings appear to lack certain global or evolutionary signals, such as long-range coevolution or organism-level priors, needed for fine-grained thermostability regression, whereas dense mutational landscapes benefit more from strong inductive biases and local reasoning. This observation suggests that scaling the PLM is not uniformly beneficial, and that architectural bias and task dynamics play a decisive role.

Finally, several experimental choices bound the scope of our conclusions. For mutational datasets (GFP, GB1), we reuse a single wild-type structure across all variants, reflecting the limited backbone changes produced by current structure predictors, like ESMFold, for near-neighbor mutations; however, this setup limits insight into cases where mutations induce genuine conformational change. We emphasize that our efficiency claims refer specifically to the number of trainable parameters and task-specific optimization cost. Structure prediction is performed once per protein as an offline preprocessing step and can be amortized across tasks, in contrast to repeated fine-tuning of large PLMs for each downstream objective. Moreover, as our pipeline relies on predicted structures, future work could explore uncertainty-aware inference to mitigate sensitivity to structural quality; especially for disordered or membrane proteins which are poorly predicted by current structure prediction methods.

## 3 Methods

### 3.1 Sequence and Structure Representations

For sequence information, we extract last-layer, per-residue embeddings from pretrained PLMs at multiple scales. Specifically, we use ESM2-T6 (8M parameters) and three sizes of ESMc: small (300M), medium (600M), and large (6B), which we denote as ESMc-S, ESMc-M, and ESMc-L, respectively. For a protein of length *n*, these embeddings form a matrix in ℝ ^*n*×*d*^, where the embedding dimension *d* is 320 for ESM2-T6 and 2560 for ESMc-L. During training, gradients from the downstream loss are not propagated back to the PLM; instead, sequence representations are extracted once and used as fixed node features.

Structural information is obtained by predicting protein 3D coordinates using ESMFold [12]. Each residue is represented by its C*α* atom, which provides a consistent and compact representation of the protein backbone. Using the C*α* coordinates, we compute an *n* × *n* pairwise distance matrix, where each entry *d*_*ij*_ denotes the Euclidean distance between residues *i* and *j*.

### 3.2 Equivariant Graph Construction

Following [35], we represent each protein as a graph

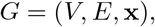

where *V* = {*v*_0_, …, *v*_*n*_} denotes the set of nodes corresponding to residues, *E* ⊆ *V* × *V* denotes the set of edges, and 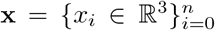 represents the 3D coordinates of the C*α* atom associated with node *v*_*i*_. Each layer of ProtEGNN takes as input the set of node embeddings 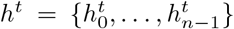, the coordinate representations 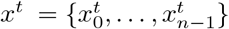, and the edge information ℰ = (*e*_*ij*_), and outputs updated node and coordinate representations *h*^*t*+1^ and *x*^*t*+1^, respectively. *t* ∈ {0, …, *T* − 1} refers to the layer number, where *T* is a hyper-parameter. Each node *v*_*i*_ is associated with an initial coordinate 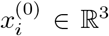, corresponding to the three-dimensional position of the C_*α*_ atom representing the residue in protein structure. Feature vectors for each node *v*_*i*_ are derived from PLM embeddings and initialized as 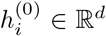. Edges are defined based on spatial proximity in 3D space using the distance matrix described above. Specifically, an edge (*v*_*i*_, *v*_*j*_) ∈ *E* exists if the Euclidean distance between the C_*α*_ atoms of residues *i* and *j* satisfies

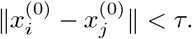

where *τ* ∈ {5, 10, 20} Angstroms (Å) and is selected as a hyper-parameter. Equivalently, we define the neighbor set 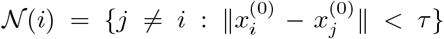 and let *E* ={(*i, j*) : *j*∈ 𝒩 (*i*)}. We optionally allow edge attributes *e*_*ij*_; in our experiments we set *e*_*ij*_ = ∅.

The layerwise equivariant message-passing is defined by the following equations:

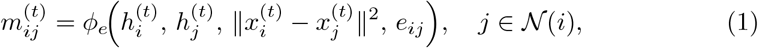

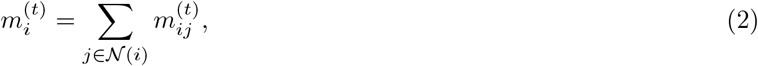

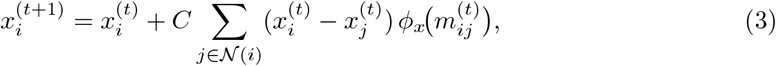

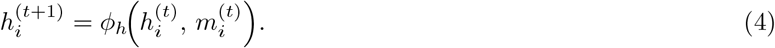

Here *ϕ*_*e*_, *ϕ*_*x*_, *ϕ*_*h*_ are edge, coordinate and node operations respectively, implemented as learnable functions using MLPs. Using relative squared distance 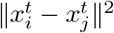 in the edge function *ϕ*_*e*_ provides rotation and translation-invariant geometric inputs and the aggregation in 2 preserves permutation invariance. 3 is the crucial coordinate update step: coordinates are moved by a weighted radial vector field where each neighbor contributes the relative displacement (*x*_*i*_ − *x*_*j*_) weighted by a learned scaler *ϕ*_*x*_(*m*_*ij*_). The constant *C* normalizes the update to stabilize magnitudes. 4 performs a permutation-equivariant update of node features using the aggregated messages 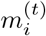, allowing feature representations to be refined in each layer.

### 3.3 Datasets

We evaluate ProtEGNN on eight datasets spanning three protein task regimes: protein property prediction, mutational-effect prediction, and protein function annotation. Together, these tasks probe global and local sequence–structure relationships under both sequence-diverse and mutation-centric settings making them a comprehensive testbed for assessing the value of explicit 3D geometry. For the ESM2-T6 backbone, pretrained and fine-tuned results in Table 1 (except for solubility) are reported from [27]. For ESMc, fine-tuning is performed using LoRA-based adaptation [50], while the pretrained baseline corresponds to frozen embeddings without task-specific updates. The second solubility dataset, Price [39], is used as a holdout set to test generalization and hence, not included in this experiment. We trained separate models for each variant on each task. All ProtEGNN models are evaluated using 3-fold cross-validation over random seeds 97, 98, 99; except solubility which used 5-fold cross-validation following [2].

#### 3.3.1 Protein Property Prediction

To assess the prediction of intrinsic protein properties that depend on global structure and physicochemical context, we evaluate solubility and stability.

##### Solubility

We use the PSI Biology solubility dataset [37], curated and cleaned by [2], containing 11,226 *E. coli* –expressed proteins labeled as soluble or insoluble. We follow the standard five-fold, label-balanced cross-validation protocol with sequence identity capped at 25%. To evaluate out-of-distribution generalization, we additionally test on the independent Price dataset [39] which contains 1323 highly expressed proteins. This dataset also does not share any sequences with identity greater than 25% to the PSI Biology dataset, ensured using USEARCH [51]. Solubility is a graph-level binary classification task and we evaluate performance using ROC AUC as the metric.

##### Stability

We evaluate protein stability using two complementary datasets that probe distinct notions of thermostability. The first is the Meltome Atlas dataset (referred to as Meltome in results), which contains ~ 23,300 protein sequences with melting temperatures (*T*_*m*_) measured via mass spectrometry. *T*_*m*_ provides a continuous-valued proxy for intrinsic thermostability and reflects both local packing interactions and long-range structural organization. We follow the train-test splits from the FLIP benchmark [41], which enforces redundancy reduction by clustering sequences at 20% pairwise sequence identity. Predictions on Meltome are formulated as a graph-level regression task and evaluated using Spearman’s rho (SPR).

The second dataset, introduced by Rocklin et al. [42] (referred to as Stability in results) from the TAPE [44] benchmark containing ~69,000 records, measures stability indirectly through protease resistance of de novo–designed mini-proteins, capturing relative fitness within local mutational neighborhoods rather than absolute thermodynamic stability. We adopt the standard TAPE split, where training and validation sets are drawn from four rounds of experimental design, and the test set consists of single-point mutants (Hamming distance 1) around 17 selected high-performing designs. This task is formulated as a graph-level regression problem analogous to learning mutational fitness landscapes evaluated using SPR.

#### 3.3.2 Mutational Effect Prediction

Fine-tuned PLMs have been shown to accurately report on mutational effects, as they implicitly capture evolutionary constraints and co-variation patterns that correlate strongly with fitness changes [52]. Given this strong baseline, our aim was to test whether explicit geometric modeling with EGNNs can be competitive in this regime. In our setup, we provide the ProtEGNN with a single wild-type structure and vary only the residue embeddings for each mutant sequence, keeping the geometry fixed. This reflects a realistic experimental scenario, where typically only one high-quality PDB structure is available and variant-specific structures are either unavailable or indistinguishable due to the limited sensitivity of structure predictors to single or few mutations.

##### GFP

From the TAPE benchmark [44], we use the fluorescence dataset (GFP), a regression task where ~ 54,000 sequences are mapped to their log-fluorescence intensity. Training variants are within Hamming distance 3 of the wild type, while test variants contain four or more mutations, explicitly probing generalization to unseen mutation combinations. GFP is a node-level regression task measured by SPR.

##### GB1

We evaluate on the GB1 landscape from the FLIP benchmark [41], which measures binding affinity of ~ 8,700 variants of the immunoglobulin-binding domain of Protein G. Mutations are introduced at four positions, producing a highly epistatic landscape. We use the standard 3-vs-rest split, where single, double, and triple mutants are used for training and more distant variants for testing. GB1 is a node-level regression task measured by SPR.

#### 3.3.3 Stable-but-Inactive variants prediction

Building on prior work [47], we employ the 11 protein systems from ProteinGym [53] evaluated by [48]. We collected multidimensional DMS measurements for single amino acid substitutions, requiring each protein to have both a functional readout and a cellular abundance readout. The resulting dataset comprised 43,725 variants in total. We trained ProtEGNN on 9,309 variants from three protein systems and evaluated generalization on 34,416 variants from the remaining eight test proteins.

### 3.4 Protein Function Annotation

#### Subcellular Localization

To evaluate functional annotation, we use the harder version of DeepLoc dataset [49], SetHard, developed by [10]. This dataset, 11,671 records, frames subcellular localization as a 10-class per-protein classification task. Sequences are filtered to remove redundancy above 20% identity between splits using MMseqs2, ensuring that predictions require generalization beyond close homologs. Subcellular localization (referred to as Sub-loc in results) is a graph-level multi-classification task measured by Accuracy (ACC).

## 4 Supplementary information

### 4.1 Robustness to alternative structure predictors

To assess the sensitivity of our pipeline to the choice of upstream structure prediction model, we performed an additional analysis in which we regenerated graphs using OpenFold3 [54] predictions in place of ESMFold structures for a subset of the solubility dataset (8,257 of the 11,225 proteins). We then re-evaluated the ESMc-M baseline using the same trained weights as in the solubility experiment in Table 1. Performance remained similar when using OpenFold3-derived graphs compared with the original ESMFold-derived graphs (0.772 and 0.769, respectively), suggesting that ProtEGNN is not strongly sensitive to the particular structure predictor used in this setting.

This is expected given the graph construction used in ProtEGNN (Section 3.1), where each protein is represented as a residue-level graph with nodes defined by C*α* coordinates and edges drawn between residues whose Euclidean distance falls below a threshold of 5, 10, or 20 Å. As the graphs depend primarily on coarse spatial proximity, relatively small coordinate differences between ESMFold and OpenFold3 predictions will often leave the graph structure largely unchanged. This explains why downstream performance remains similar: for the vast majority of proteins in this benchmark, replacing ESMFold with OpenFold3 does not substantially alter the geometric information presented to the model.

### 4.2 Hyperparamters

For completeness and reproducibility, we provide the key hyperparameters used in ProtEGNN in Table 6. The table lists the default values used in our experiments, the search space considered during tuning, and a brief description of the role of each parameter. The exact configuration files and hyperparameter settings used for each model are available in our GitHub repository.

**Table 6.**
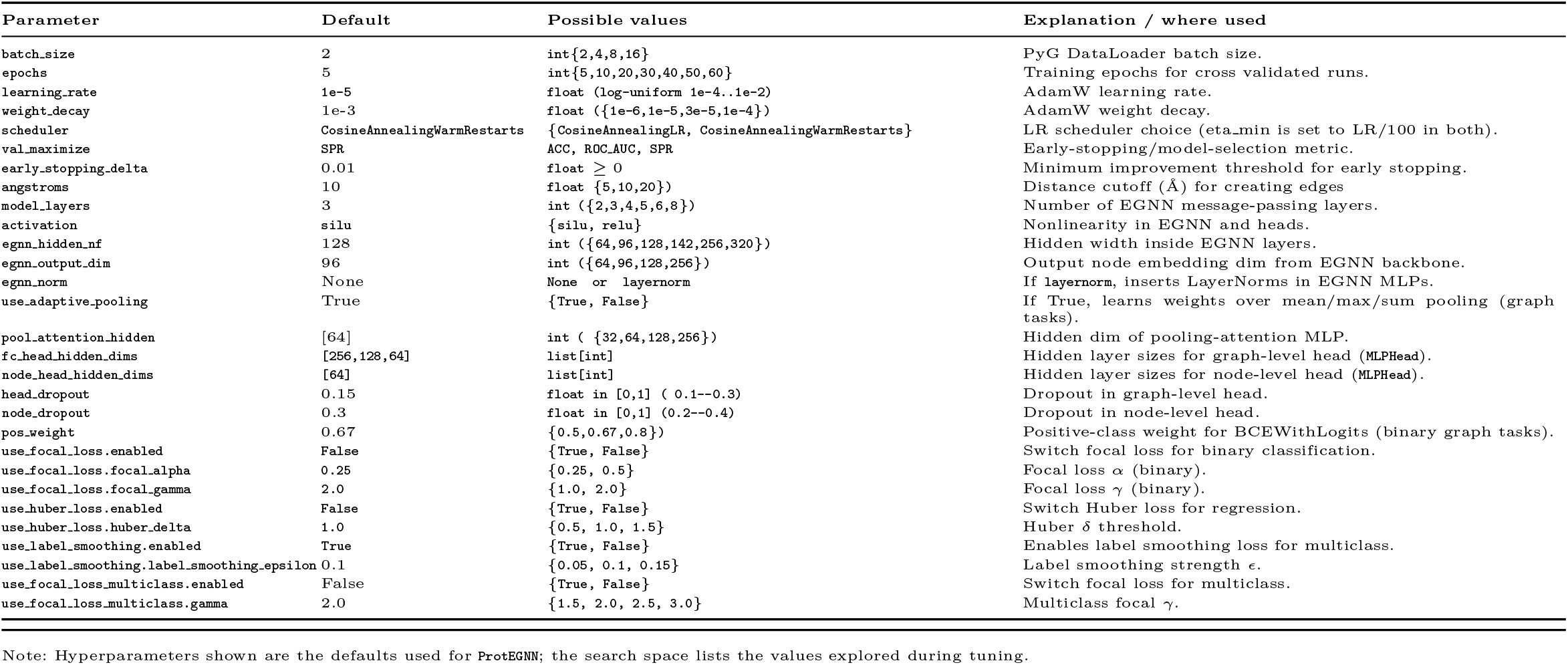
Hyperparameters in ProtEGNN.

## References

[1] Tsuboyama, K., Dauparas, J., Chen, J., Laine, E., Mohseni Behbahani, Y., Weinstein, J.J., Mangan, N.M., Ovchinnikov, S., Rocklin, G.J.: Mega-scale experimental analysis of protein folding stability in biology and design. Nature 620(7973), 434–444 (2023) 10.1038/s41586-023-06328-6

[2] Thumuluri, V., Martiny, H.-M., Almagro Armenteros, J.J., Salomon, J., Nielsen, H., Johansen, A.R.: Netsolp: predicting protein solubility in escherichia coli using language models. Bioinformatics 38(4), 941–946 (2021) 10.1093/bioinformatics/btab801

[3] Yu, H., Deng, H., He, J., Keasling, J.D., Luo, X.: Unikp: a unified framework for the prediction of enzyme kinetic parameters. Nature communications 14(1), 8211 (2023)

[4] Liu, Q., Xun, G., Feng, Y.: The state-of-the-art strategies of protein engineering for enzyme stabilization. Biotechnology Advances 37(4), 530–537 (2019) 10.1016/j.biotechadv.2018.10.011

[5] Sellés Vidal, L., Isalan, M., Heap, J.T., Ledesma-Amaro, R.: A primer to directed evolution: current methodologies and future directions. RSC Chemical Biology 4(4), 271–291 (2023) 10.1039/d2cb00231k

[6] Watson, J.L., Juergens, D., Bennett, N.R., Trippe, B.L., Yim, J., Eisenach, H.E., Ahern, W., Borst, A.J., Ragotte, R.J., Milles, L.F., et al.: De novo design of protein structure and function with rfdiffusion. Nature 620(7976), 1089–1100 (2023)

[7] Butcher, J., Krishna, R., Mitra, R., Brent, R.I., Li, Y., Corley, N., Kim, P.T., Funk, J., Mathis, S., Salike, S., et al.: De novo design of all-atom biomolecular interactions with rfdiffusion3. bioRxiv (2025)

[8] Stark, H., Faltings, F., Choi, M., Xie, Y., Hur, E., O’Donnell, T., Bushuiev, A., Uçar, T., Passaro, S., Mao, W., et al.: Boltzgen: Toward universal binder design. bioRxiv, 2025–11 (2025)

[9] Weissenow, K., Heinzinger, M., Rost, B.: Protein language-model embeddings for fast, accurate, and alignment-free protein structure prediction. Structure 30(8), 1169–1177 (2022)

[10] Stärk, H., Dallago, C., Heinzinger, M., Rost, B.: Light attention predicts protein location from the language of life. Bioinformatics Advances 1(1), 035 (2021)

[11] Marquet, C., Heinzinger, M., Olenyi, T., Dallago, C., Erckert, K., Bernhofer, M., Nechaev, D., Rost, B.: Embeddings from protein language models predict conservation and variant effects. Human genetics 141(10), 1629–1647 (2022)

[12] Lin, Z., Akin, H., Rao, R., Hie, B., Zhu, Z., Lu, W., Smetanin, N., Verkuil, R., Kabeli, O., Shmueli, Y., Santos Costa, A., Fazel-Zarandi, M., Sercu, T., Candido, S., Rives, A.: Evolutionary-scale prediction of atomic level protein structure with a language model. Cold Spring Harbor Laboratory (2022). 10.1101/2022.07.20.500902. http://dx.doi.org/10.1101/2022.07.20.500902

[13] Rives, A., Meier, J., Sercu, T., Goyal, S., Lin, Z., Liu, J., Guo, D., Ott, M., Zitnick, C.L., Ma, J., Fergus, R.: Biological structure and function emerge from scaling unsupervised learning to 250 million protein sequences. Cold Spring Harbor Laboratory (2019). 10.1101/622803. http://dx.doi.org/10.1101/622803

[14] ESM Team: ESM Cambrian: Revealing the mysteries of proteins with unsupervised learning. EvolutionaryScale Website (2024). https://evolutionaryscale.ai/blog/esm-cambrian Accessed 2024-12-04

[15] Elnaggar, A., Heinzinger, M., Dallago, C., Rehawi, G., Wang, Y., Jones, L., Gibbs, T., Feher, T., Angerer, C., Steinegger, M., Bhowmik, D., Rost, B.: Prottrans: Towards cracking the language of life’s code through self-supervised learning. Cold Spring Harbor Laboratory (2020) 10.1101/2020.07.12.199554

[16] Alsamkary, H., Elshaffei, M., Elkerdawy, M., Elnaggar, A.: Ankh3: Multi-task pretraining with sequence denoising and completion enhances protein representations. arXiv preprint arXiv:2505.20052 (2025)

[17] Olsen, T.H., Moal, I.H., Deane, C.M.: Ablang: an antibody language model for completing antibody sequences. Bioinformatics Advances 2(1), 046 (2022)

[18] Outeiral, C., Deane, C.M.: Codon language embeddings provide strong signals for use in protein engineering. Nature Machine Intelligence 6(2), 170–179 (2024)

[19] Rao, R., Meier, J., Sercu, T., Ovchinnikov, S., Rives, A.: Transformer protein language models are unsupervised structure learners. Cold Spring Harbor Laboratory (2020) 10.1101/2020.12.15.422761

[20] Vig, J., Madani, A., Varshney, L.R., Xiong, C., Socher, R., Rajani, N.F.: BERTology Meets Biology: Interpreting Attention in Protein Language Models. arXiv (2020). 10.48550/ARXIV.2006.15222. https://arxiv.org/abs/2006.15222

[21] Lombard, V., Timsit, D., Grudinin, S., Laine, E.: Seamoon: from protein language models to continuous structural heterogeneity (2024) 10.1101/2024.09.23.614585

[22] Meng, Y., Zhang, Z., Zhou, C., Tang, X., Hu, X., Tian, G., Yang, J., Yao, Y.: Protein structure prediction via deep learning: an in-depth review. Frontiers in Pharmacology 16 (2025) 10.3389/fphar.2025.1498662

[23] Abramson, J., Adler, J., Dunger, J., Evans, R., Green, T., Pritzel, A., Ronneberger, O., Willmore, L., Ballard, A.J., Bambrick, J., Bodenstein, S.W., Evans, D.A., Hung, C.-C., O’Neill, M., Reiman, D., Tunyasuvunakool, K., Wu, Z., Žemgulytė, A., Arvaniti, E., Beattie, C., Bertolli, O., Bridgland, A., Cherepanov, A., Congreve, M., Cowen-Rivers, A.I., Cowie, A., Figurnov, M., Fuchs, F.B., Gladman, H., Jain, R., Khan, Y.A., Low, C.M.R., Perlin, K., Potapenko, A., Savy, P., Singh, S., Stecula, A., Thillaisundaram, A., Tong, C., Yakneen, S., Zhong, E.D., Zielinski, M., Žídek, A., Bapst, V., Kohli, P., Jaderberg, M., Hassabis, D., Jumper, J.M.: Accurate structure prediction of biomolecular interactions with alphafold 3. Nature 630(8016), 493–500 (2024) 10.1038/s41586-024-07487-w

[24] Ferreira, J., Castro, F.: Advances in protein solubility and thermodynamics: quantification, instrumentation, and perspectives. CrystEngComm 25(46), 6388–6404 (2023) 10.1039/d3ce00757j

[25] Vendruscolo, M., Knowles, T.P.J., Dobson, C.M.: Protein solubility and protein homeostasis: A generic view of protein misfolding disorders. Cold Spring Harbor Perspectives in Biology 3(12), 010454–010454 (2011) 10.1101/cshperspect.a010454

[26] Jiang, F., Li, M., Dong, J., Yu, Y., Sun, X., Wu, B., Huang, J., Kang, L., Pei, Y., Zhang, L., Wang, S., Xu, W., Xin, J., Ouyang, W., Fan, G., Zheng, L., Tan, Y., Hu, Z., Xiong, Y., Feng, Y., Yang, G., Liu, Q., Song, J., Liu, J., Hong, L., Tan, P.: A general temperature-guided language model to design proteins of enhanced stability and activity. Science Advances 10(48) (2024) 10.1126/sciadv.adr2641

[27] Schmirler, R., Heinzinger, M., Rost, B.: Fine-tuning protein language models boosts predictions across diverse tasks. Nature Communications 15(1) (2024) 10.1038/s41467-024-51844-2

[28] Zhang, X., Hu, X., Zhang, T., Yang, L., Liu, C., Xu, N., Wang, H., Sun, W.: Plm sol: predicting protein solubility by benchmarking multiple protein language models with the updated escherichia coli protein solubility dataset. Briefings in Bioinformatics 25(5) (2024) 10.1093/bib/bbae404

[29] Yuan, Y., Luo, H., Tian, Y.: Lmprotein: a protein language model based framework for protein structural property prediction. Physical Chemistry Chemical Physics 28(2), 1747–1758 (2026) 10.1039/d5cp01861g

[30] Bikias, T., Stamkopoulos, E., Reddy, S.T.: Plmfit: benchmarking transfer learning with protein language models for protein engineering. Briefings in Bioinformatics 26(4) (2025) 10.1093/bib/bbaf381

[31] McCloskey, M., Cohen, N.J.: Catastrophic interference in connectionist networks: The sequential learning problem. In: Psychology of Learning and Motivation vol. 24, pp. 109–165. Elsevier, (1989)

[32] Heinzinger, M., Weissenow, K., Sanchez, J.G., Henkel, A., Mirdita, M., Steinegger, M., Rost, B.: Bilingual language model for protein sequence and structure. NAR Genomics and Bioinformatics 6(4) (2024) 10.1093/nargab/lqae150

[33] Su, J., Han, C., Zhou, Y., Shan, J., Zhou, X., Yuan, F.: Saprot: Protein language modeling with structure-aware vocabulary (2023) 10.1101/2023.10.01.560349

[34] Li, M., Tan, Y., Ma, X., Zhong, B., Yu, H., Zhou, Z., Ouyang, W., Zhou, B., Tan, P., Hong, L.: Prosst: Protein language modeling with quantized structure and disentangled attention. Advances in Neural Information Processing Systems 37, 35700–35726 (2024)

[35] Satorras, V.G., Hoogeboom, E., Welling, M.: E(n) Equivariant Graph Neural Networks. arXiv (2021). 10.48550/ARXIV.2102.09844. https://arxiv.org/abs/2102.09844

[36] Elnaggar, A., Essam, H., Salah-Eldin, W., Moustafa, W., Elkerdawy, M., Rochereau, C., Rost, B.: Ankh: Optimized protein language model unlocks general-purpose modelling. arXiv preprint arXiv:2301.06568 (2023)

[37] Bhandari, B.K., Gardner, P.P., Lim, C.S.: Solubility-weighted index: fast and accurate prediction of protein solubility. Bioinformatics 36(18), 4691–4698 (2020) 10.1093/bioinformatics/btaa578

[38] Rao, R.M., Liu, J., Verkuil, R., Meier, J., Canny, J., Abbeel, P., Sercu, T., Rives, A.: Msa transformer. In: International Conference on Machine Learning, pp. 8844–8856 (2021). PMLR

[39] Price, W.N., Handelman, S.K., Everett, J.K., Tong, S.N., Bracic, A., Luff, J.D., Naumov, V., Acton, T., Manor, P., Xiao, R., Rost, B., Montelione, G.T., Hunt, J.F.: Large-scale experimental studies show unexpected amino acid effects on protein expression and solubility in vivo in e. coli. Microbial Informatics and Experimentation 1(1) (2011) 10.1186/2042-5783-1-6

[40] Tan, Y., Zheng, J., Hong, L., Zhou, B.: Protsolm: Protein solubility prediction with multi-modal features. In: 2024 IEEE International Conference on Bioinformatics and Biomedicine (BIBM), pp. 223–232 (2024). IEEE

[41] Dallago, C., Mou, J., Johnston, K.E., Wittmann, B.J., Bhattacharya, N., Goldman, S., Madani, A., Yang, K.K.: Flip: Benchmark tasks in fitness landscape inference for proteins. “ (2021) 10.1101/2021.11.09.467890

[42] Rocklin, G.J., Chidyausiku, T.M., Goreshnik, I., Ford, A., Houliston, S., Lemak, A., Carter, L., Ravichandran, R., Mulligan, V.K., Chevalier, A., Arrowsmith, C.H., Baker, D.: Global analysis of protein folding using massively parallel design, synthesis, and testing. Science 357(6347), 168–175 (2017) 10.1126/science.aan0693

[43] Yang, K.K., Fusi, N., Lu, A.X.: Convolutions are competitive with transformers for protein sequence pretraining. Cell Systems 15(3), 286–294 (2024)

[44] Rao, R., Bhattacharya, N., Thomas, N., Duan, Y., Chen, P., Canny, J., Abbeel, P., Song, Y.: Evaluating protein transfer learning with tape. Advances in neural information processing systems 32 (2019)

[45] Wang, Z., Combs, S.A., Brand, R., Calvo, M.R., Xu, P., Price, G., Golovach, N., Salawu, E.O., Wise, C.J., Ponnapalli, S.P., Clark, P.M.: Lm-gvp: an extensible sequence and structure informed deep learning framework for protein property prediction. Scientific Reports 12(1) (2022) 10.1038/s41598-022-10775-y

[46] Hsu, C., Verkuil, R., Liu, J., Lin, Z., Hie, B., Sercu, T., Lerer, A., Rives, A.: Learning inverse folding from millions of predicted structures. Cold Spring Harbor Laboratory (2022). 10.1101/2022.04.10.487779. http://dx.doi.org/10.1101/2022.04.10.487779

[47] Cagiada, M., Jonsson, N., Lindorff-Larsen, K.: Decoding molecular mechanisms for loss-of-function variants in the human proteome (2024) 10.1101/2024.05.21.595203

[48] Ding, K., Li, Z., Tu, T., Luo, J., Luo, Y.: Deconvolving mutation effects on protein stability and function with disentangled protein language models (2026) 10.64898/2026.02.03.703560

[49] Almagro Armenteros, J.J., Sønderby, C.K., Sønderby, S.K., Nielsen, H., Winther, O.: Deeploc: prediction of protein subcellular localization using deep learning. Bioinformatics 33(21), 3387–3395 (2017)

[50] Hu, E.J., Shen, Y., Wallis, P., Allen-Zhu, Z., Li, Y., Wang, S., Wang, L., Chen, W., et al.: Lora: Low-rank adaptation of large language models. ICLR 1(2), 3 (2022)

[51] Edgar, R.C.: Search and clustering orders of magnitude faster than blast. Bioinformatics 26(19), 2460–2461 (2010) 10.1093/bioinformatics/btq461

[52] Glaser, M., Brägelmann, J.: Esm-effect: An effective and efficient fine-tuning framework towards accurate prediction of mutation’s functional effect (2025) 10.1101/2025.02.03.635741

[53] Notin, P., Kollasch, A.W., Ritter, D., Niekerk, L., Paul, S., Spinner, H., Rollins, N., Shaw, A., Weitzman, R., Frazer, J., Dias, M., Franceschi, D., Orenbuch, R., Gal, Y., Marks, D.S.: Proteingym: Large-scale benchmarks for protein design and fitness prediction (2023) 10.1101/2023.12.07.570727

[54] The OpenFold3 Team: OpenFold3-preview. 10.5281/zenodo.19001000. https://github.com/aqlaboratory/openfold-3

